# Negative allometry of egg size among 29 species of drosophilid flies

**DOI:** 10.64898/2026.03.15.711943

**Authors:** Jonathan A. Rader, Mary Petersen, Dariel A. Cortés, Daniel R. Matute

## Abstract

The body size of adults and immature stages are fundamental animal traits that influence animal physiology, ecology, and range distribution. While the importance of egg size has been acknowledged as a proxy of parental investment in animals, little work has addressed the tempo and mode of evolution of egg size and shape. Here, we present a comparative study of this trait using a phylogeny based on genome-wide markers together with measurements of egg size and adult body size from 29 drosophilid species. Our analyses revisit the allometric relationship between egg size and body size and show that egg size scales negatively with respect to adult size, even after accounting for shared evolutionary history. In other words, larger species tend to produce proportionally smaller eggs. We also detect a moderate phylogenetic signal in both egg size and egg shape, indicating that closely related species resemble each other in these traits. Model comparisons show that the evolution of egg morphology in drosophilids is best described by gradual divergence through time driven by stochastic evolutionary change. This pattern contrasts with findings from other animal groups, including birds, cephalopods, and reptiles, where alternative evolutionary models better explain trait evolution. Together, these results suggest that the evolutionary dynamics shaping egg morphology in drosophilids differ from those operating in other major lineages and underscore the importance of comparative analyses of early developmental traits across taxa.

## Introduction

Body size is a fundamental animal trait that influences an organism’s physiology, ecology, and evolutionary dynamics (Roff, 1986; Klingenberg & Spence, 1997). Many life-history traits are tightly correlated with body size. Lifespan (Lindstedt & Calder, 1981; Speakman, 2005), development time (Blueweiss *et al*., 1978; Nijhout *et al*., 2010), as well as overall growth and survival (Schmidt-Nielsen, 1984) are all correlated with body size. Larger organisms tend to have longer lifespans and delayed reproduction, whereas smaller organisms often reproduce at earlier ages and in greater numbers (Peters, 1983). Body size also affects physiological processes such as the rate of diffusion of nutrients and waste disposal (Schmidt-Nielsen, 1984; Blanckenhorn, 2000; Gokhale & Shingleton, 2015), (Schmidt-Nielsen, 1984). Ecological features such as range size (Gaston & Blackburn, 1996), predation and competition (Brown, 1984), and dispersal ability (Jenkins *et al*., 2007; De Bie *et al*., 2012) also depend on body size. Body size is also correlated with mitochondrial substitution rate (Fontanillas *et al*., 2007), and diversification rates (Amado *et al*., 2021), but see (Feldman *et al*., 2016; Lee *et al*., 2016).

Egg size is associated with critical life-history traits, including adult body size and varies with adult body size (García-Barros, 2000; Yanagi & Tuda, 2012) but see (Church *et al*., 2019), development time (Berrill, 1935; Levitan, 2000), and fecundity (Steele & Steele, 2020). Egg size, a proxy of parental investment (Berrigan, 1991; Fox & Czesak, 2000), scales positively with respect to adult body size across a wide range of taxa, including birds (Olsen *et al*., 1994; Rotenberry & Balasubramaniam, 2020), amphibians (Hallmann & Griebeler, 2020; Cejp & Griebeler, 2024), reptiles (Hallmann & Griebeler, 2018), and insects (Berrigan, 1991; García-Barros, 2000). In seven insect orders, adult body size and egg volume show a negative allometric relationship, but the predictive power of the scaling relationships varies widely across clades (Church *et al*., 2019). Egg size has been hypothesized to predict development duration (Maino *et al*., 2017), although the strength and predictive power of this scaling vary among clades (Church *et al*., 2019). Despite this body of work, relatively little effort has been devoted to evaluating the evolutionary dynamics of immature-stage morphology, including eggs and embryos. One notable exception concerns egg size and shape, both of which show strong phylogenetic signal in birds (Shatkovska *et al*., 2018). Thus, even with extensive data compilations (Berrigan, 1991; Hendriks & Mulder, 2008; Church *et al*., 2019), a systematic appraisal of the tempo and mode of evolution of egg morphology in any insect – while accounting for phylogenetic and geographic history – is lacking.

*Drosophila* has long provided a model system for the evolutionary study of morphology (reviewed in (Markow, 2015), ecology (e.g., (Spieth & Heed, 1972; Markow & O’Grady, 2008; Mendes *et al*., 2021), and the genetic basis of those traits (e.g., Barker *et al*., 2013). In particular, studies in *Drosophila* have revealed the relative contributions of environmental conditions and genetic components to body size (Robertson, 1960, 1966; Partridge *et al*., 1994; Zamudio *et al*., 1995). The genus harbors and contains two large radiations (the *Drosophila* and *Sophophora* subgenus) that diverged approximately 50 million years ago (Tamura *et al*., 2004; Russo *et al*., 2013; Suvorov *et al*., 2022). *Drosophila*’s short gen time, experimental tractability, and the abundance of genomic data make *Drosophila* a natural candidate to test evolutionary hypotheses. Previously, this opportunity was leveraged to study the presumed tradeoff between egg size and developmental time; a survey of 11 species revealed differences among species in egg size and larval eclosion time but no correlation between egg size and developmental time (Markow *et al*., 2009). Since then, we and others (Suvorov *et al*., 2022; Kim *et al*., 2024) built a time-calibrated tree based on genome-wide markers, which enables a more comprehensive and detailed study of the macroevolutionary patterns in the genus *Drosophila*.

Thus, the time is ripe for a phylogenetically-informed assessment of egg size evolution in *Drosophila*. In this piece, we provide the first such assessment using a time-calibrated phylogenetic tree and egg and body size measurements for 29 drosophilid species. We provide a new assessment of allometric scaling and confirm that egg size and adult body size have a negative allometric relationship, even after correcting for phylogenetic relationships; larger fly species produce comparatively small eggs. We also find moderate phylogenetic signal in egg size and shape variation. The evolution of these two traits is best explained by a Brownian motion model, in which the trait means change through time according to a random drift process. This pattern suggests that egg evolution in drosophilids differs from that observed in birds, cephalopods, and reptiles, for which Brownian motion does not provide the best fit.

Overall, our results indicate that the process of evolution of egg morphology in *Drosophila* differs from the process in other lineages, highlighting the need for comparative analyses of developmental stages across taxa.

## Methods and Materials

### Fly stocks and virgin collection

We collected and measured egg size for 70 lines of 29 species of drosophilid flies (Table S1). Flies were either collected from the field or purchased at the Cornell Stock Center. Lines were derived from a single field-collected inseminated female and maintained as a shelf-stable stock (isofemale lines). All lines were maintained in an incubator with a consistent 12-hour light/dark cycle at 24°C during photophase and 21°C during scotophase. The metadata for each species is also listed in Table S1. To collect eggs, we expanded each isofemale line for the study from 30 mL vials into 100 mL plastic bottles with cornmeal and yeast (35 g/L, Red Star active dry yeast, *Saccharomyces cerevisiae*). We added a pupation substrate (Kimwipes Delicate Task; Kimberly Clark, Irving, TX) to facilitate pupation. We cleared each bottle daily before the onset of photophase. Males and females were separated as virgins and kept in groups of up to 25 individuals in 30mL cornmeal plastic vials.

After four days, we mixed females and males (20-25 individuals per sex) in an empty oviposition cage (Flystuff, San Diego, CA, USA) at 24°C with a yeast paste and wet paper to avoid desiccation. After 12 hours, we provided an oviposition substrate, apple juice agar. From these cages, we collected freshly deposited eggs (less than six hours after oviposition). We changed the agar plate every 24 hours to collect and measure eggs minimizing larval eclosion. We repeated the same approach for three days, at which point we collected the flies and weighed them in separate batches of males and females (n∼20) on a Mettler-Toledo AB54-S balance (precision: 0.1 mg, resolution: 0.1 mg; Mettler-Toledo, Columbus, OH). By weighing the flies in batches and dividing by the count of the flies, we obtained an average mass for the male and female flies. (Dead flies were not included in this estimate). We note that we were unable to take individual mass measurements for each fly, and therefore relied on average masses in subsequent analyses.

### Egg measurements

We used image analysis to measure fly egg size. Eggs were harvested from the egg collection chambers (average *n* = 686 eggs per species) and placed onto fresh agar in a petri dish alongside a 1mm graduated scale bar. We photographed the eggs using a Jenoptix Gryphax microscope camera (Arktur 8MP SKU: 571025) affixed to a Leica dissecting microscope at 20X magnification. Each image was processed to isolate and measure the areas of each egg using a custom program in Fiji (Schindelin *et al*., 2012), which is an open-source implementation of ImageJ (Rashband, 1997; Abramoff *et al*., 2004). We thresholded images and used a blob-detection routine to isolate eggs from the image background. We measured the pixel area of each egg in the images and converted them to mm^2^ using the scale bar for each image.

We calculated two metrics of egg morphology. First, we calculated egg area as an ellipse using the formula:

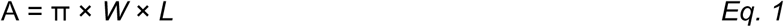

where *L* is the length (longer axis), and *W* is the width (shorter axis). Previous surveys have calculated the volume of the egg on the assumption that depth is equivalent to W, but we restrict our analyses to egg area to minimize assumptions. Second, we calculated the egg aspect ratio as L/W.

### Phylogenetic tree

We used the phylogenetic reconstruction of drosophilid flies from (Kim *et al*., 2021, 2024; Suvorov *et al*., 2022) for our phylogenetic analyses. Briefly, this maximum likelihood tree is based on whole genomes of over 300 species of drosophilids and was time-calibrated with fossil records. Because subsequent analyses (e.g., evolutionary rate analysis and ancestral character state reconstruction via maximum likelihood) require an ultrametric tree structure, we used the penalized likelihood method from Sanderson (2002).We scaled the tree to relative time using the *makeChronosCalib* function to create time calibration points and then fitting a phylogenetic model to those points using the *‘chronos’* function (both functions are in the *ape* R package; Paradis *et al*., 2004), while allowing for rate variation across the tree (model = “relaxed”). We tested multiple molecular clock model fits with varying values of L (ranging from 0 to 10), and evaluated the best fit using the Penalized Hierarchical Information Criterion (PHIC). We pruned the resulting tree (without altering branch lengths) using the *drop.tip* function in the *ape* package (Paradis *et al*., 2004) to include only the species in our egg dataset (See Figure 1).

**FIGURE 1.**
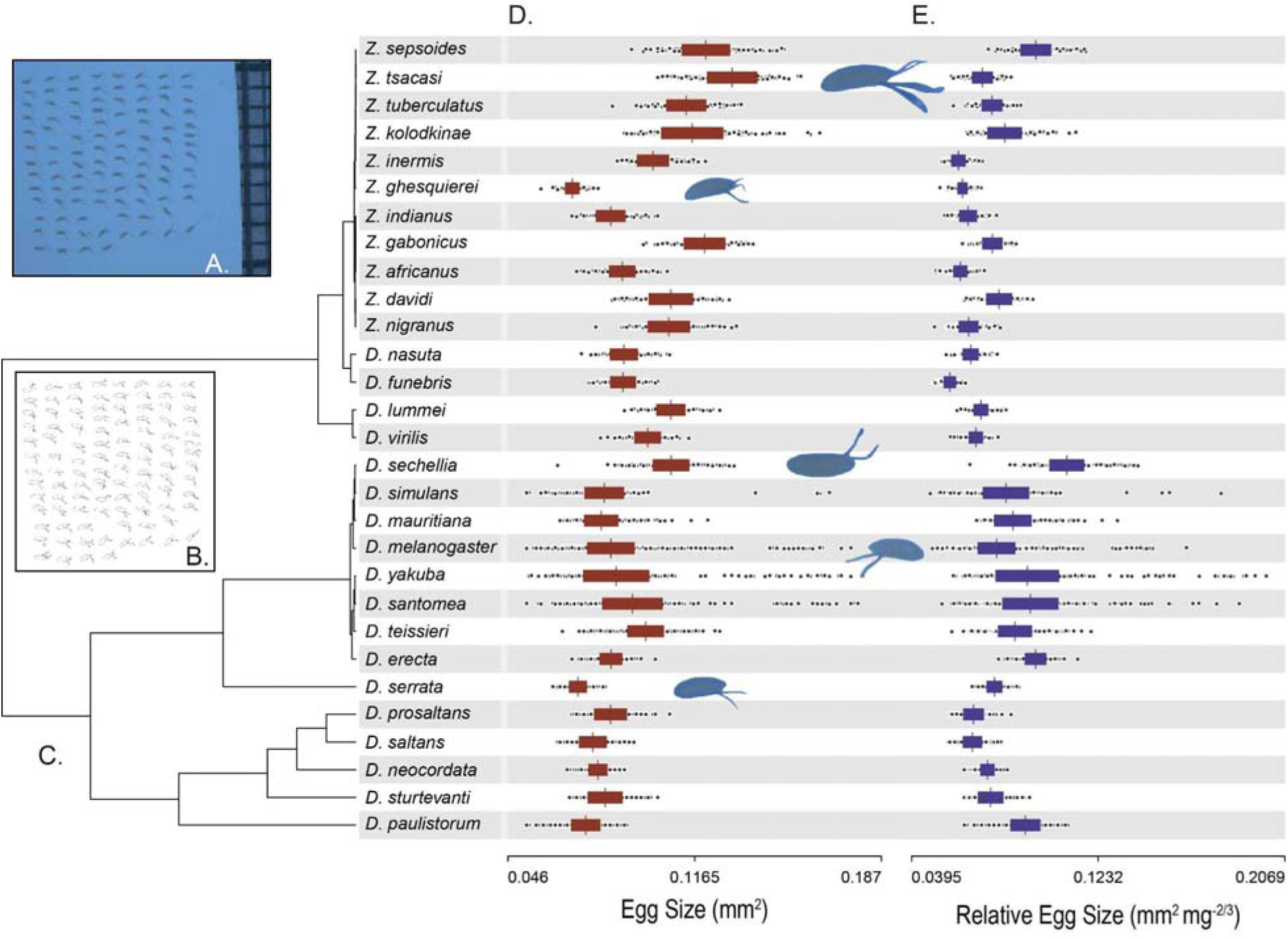
Image analysis workflow, phylogenetic tree, and variation of egg size across the phylogeny. **A.** An example of a photograph of eggs. **B.** Output of the image acquisition routine. **C.** Maximum likelihood phylogenetic tree of the 29 species of drosophilids included in this study. **D.** Egg size distribution. We include four representative examples of egg size. **E.** Egg size normalized by adult body mass.

### Tests for allometric and isometric scaling

Allometric scaling relationships typically take the form of a power function:

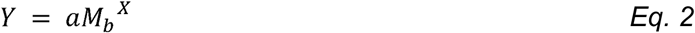

where *Y* is a trait that varies with body mass (*M_b_*), *a* is a taxon-specific constant, and *X* is the scaling factor that describes the change in *Y* with respect to *M_b_*. A log_10_ transformation of Eq. 1 linearizes the power function, rendering the exponential scaling factor *X* as the slope in the transformed equation:

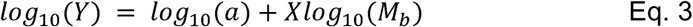

Under an assumption of isometry, wherein a trait maintains exact geometric similarity among taxa across a range of body sizes, we expect a scaling factor of *X*=1. This also assumes that the dimensions of the two traits being compared are equal; unequal dimensions yield different isometric expectations. For example, a trait measured in units of area (such as wing area) vs. a trait that is proportional to volume (like *M_b_*) would yield an isometric expectation of *X*=0.67. Egg volume or mass would be an ideal measurement, but is complicated to record. Our measurement of egg size is two-dimensional; we measured size as area (units: *length*^2^) in photographs. This decision created a dimensional mismatch between our egg size proxy and the measurements of adult body size (average body mass of adult flies, *M_b_*). Mass is proportional to volume, or *length*^3^. We conducted analyses of the scaling relationship between eggs and adult size on log_10_ transformations of both trait values. The null expectation of isometric scaling would thus yield a slope of 0.67. Slopes > 0.67 would indicate positive allometry, where larger-bodied flies have disproportionately large eggs, and slopes < 0.67 would show negative allometry and disproportionately small eggs in larger-bodied flies.

We used Phylogenetic Generalized Least Squares (PGLS) analyses (Grafen, 1989; Martins & Hansen, 1997; Pagel, 1999) implemented with the *pgls* function in the *caper* R package (Orme *et al*., 2018) to estimate the relationship between egg size and the average body mass of adult female flies. We ran separate PGLS models with maximum-likelihood optimization (Freckleton *et al*., 2002) for each of three branch length transformations, lambda (□), kappa (□), and delta (□). These branch length transformations provide complementary insight into the effects of phylogenetic relatedness on the relationships between traits. A □ transformation investigates the influence of tree topology by raising the branch lengths to the power of □. When □ = 1 there is no alteration to the branch lengths, but □ = 0 forces all branch lengths to equal 1 (O’Meara, 2012). Delta transformations stretch either early (□ < 1) or later (□ > 1) branches, demonstrating whether trait evolution occurred early and deep within the tree, or more recently near the species tips, respectively (O’Meara, 2012). We ran these PGLS analyses on the entire species-level phylogeny, and we iterated the analyses within each of the subgenera. Comparison of these PGLS models allowed us to assess whether the species-level pattern 1) held within species and across genera, and 2) if our results were robust to varying assumptions about the influence of tree topology.

### Phylogenetic signal

We tested for patterns of Phylogenetic Conservatism (PC) in egg size and adult body size with two complementary metrics of phylogenetic signal, Blomberg’s *K* (Blomberg *et al*., 2003) and Pagel’s λ (Pagel, 1999). The *K* statistic compares the similarity among relatives with that expected from a Brownian motion (BM) model of evolutionary change along the branches of a phylogenetic tree to determine if the trait value is more or less divergent than would be predicted by random evolution (Blomberg *et al*., 2003). When *K* equals 1, there is a close correspondence between observed phenotypic values and the BM null prediction. When *K* < 1, trait values vary more than would be predicted by BM evolution, and is evidence against a PC pattern (Losos, 2008; Wiens *et al*., 2010; Cooper *et al*., 2011). If *K* is larger than 1, trait values are more similar to each other than BM evolution would predict, indicating a pattern of PC, but not confirming PC (though this may be insufficient to confirm PC, see Wiens *et al*., 2010).

Pagel’s λ (Pagel, 1999) is a measure of the extent to which the phylogenetic history of a clade is predictive of the trait distribution at the tree tips. The λ statistic ranges between 0 and 1; when λ = 0, the phylogenetic structure is the equivalent of a star phylogeny, with all tips emerging from a single node with equal branch lengths. When λ = 1, the phylogenetic structure conforms to the BM expectation as determined by tree topology and branch lengths. We used the ‘*phylosig*’ function in the *phytools* R package (Revell, 2012) to calculate both K and λ, with 1,000 simulations to determine if the calculated value differed from zero. We tested the significance of the calculated *K* values recovered for the phylogeny to those for an iterative tip-shuffling randomization (Kembel *et al*., 2010). The significance of lambda was evaluated using a likelihood ratio test (Revell & Revell, 2014). We additionally iterated the λ phylogenetic signal tests for egg size for the two subgenera in our sample, *Sophophora* and *Drosophila*.

### Models of trait evolution and evolutionary rate estimates

We identified the best-fitting model of trait evolution of egg and adult body size. We fit six different models of phenotypic evolution (BM, Ornstein-Uhlenbeck–OU–, Kappa, Early Burst–EB–, Rate Trend–RT–, and Delta) and one non-phylogenetic model (White noise, WN) to both egg size and adult body size with the *fitContinuous* function in the *geiger* R package (Harmon *et al*., 2008; Pennell *et al*., 2014). We used AICc to identify the best-fit evolutionary model for each trait. Table S2 describes the specifics and parameters of each of the seven fitted models. We repeated these model fits for egg size in each of the individual clades, but with a more restricted set of candidate models: BM, OU, and the non-evolutionary WN model.

We then conducted an ancestral character state reconstruction to look for significant changes in trait evolution rate in egg size among drosophilids. We used a maximum likelihood approach to estimate ancestral body size with the function *anc.ML* in the *phytools* package (Revell, 2012; Revell & Revell, 2014) under both BM and OU regimes. We considered evolutionary transitions to be significant if the 95% confidence intervals of descendant node values did not overlap with those of their ancestors. This provides a conservative estimate of significant differentiation of mean between adjacent tree nodes and/or tips. Overlapping confidence intervals are consistent with gradual evolutionary change; non-overlapping intervals are indicative of a rapid divergence event.

### Evolutionary rate of traits and multirate comparison

We investigated the pace of morphological evolution in egg size and adult body mass. We calculated the evolutionary rate parameter (□^2^ - the rate of change in trait mean under a BM process) using the *fitcontinuous* function as described above. We used the wAIC values to compare support for the evolutionary rate estimates generated by each of the evolutionary models. We also evaluated whether the trait evolutionary rates were similar across the *Drosophila* phylogeny, or whether they varied throughout the history of the lineage. To do this, we use maximum-likelihood models of evolutionary rate of egg and adult size with the *multirateBM* function in (R *phytools*, Revell & Revell, 2014). These models use a penalized likelihood approach to fit a flexible BM model (Revell, 2021), which allows for rate shifts along the tree. To facilitate comparison of the resulting patterns among traits, we rescaled the σ^2^ output from each *multirateBM* model to a range of 0-1 by dividing branch rate values by the maximum rate for a given trait. We next calculated a difference index by subtracting the evolutionary rates of egg size from those of body mass. This index spans from −1 to 1, where a value of 0 indicates that both traits evolve at the same rate. Positive values reflect a slower evolution of egg size relative to body mass, whereas negative values indicate comparatively faster evolution of egg size relative to mass.

## RESULTS

### Egg size differs between species

We measured 19,895 eggs from 70 genotypes representing 29 species of drosophilid flies in two genera, *Drosophila* and *Zaprionus* (Figure 1; Note that *Zaprionus* is nested within the *Drosophila* subgenus; (Yassin *et al*., 2008, 2010; Suvorov *et al*., 2022). Mean egg size (±1 sd) across all species was 0.090 ± 0.016 mm^2^. We found significant differences across species (Figure 1, Figure S1, *F*_28,19886_= 1543.7, *p* < 1 × 10^-10^) and over 90% of the pairwise comparisons are significant (Table S3). Egg size ranged from 0.065 ± 0.003 mm^2^ for *Zaprionus ghesquierei* to 0.132 ± 0.011 mm^2^ (*t* = -21.684, *p* < 1 × 10^-10^) for *Z. tsacasi* (*t* = 5.080×10^-2^, *p* < 1 × 10^-10^; Figure 1). This twofold difference is slightly larger than the previous estimates of egg size (70% area’s difference between *D. sechellia* and *D. pseudoobscura*, Table 1 in (Markow *et al*., 2009).

**TABLE 1.** Isofemale lines of the same species show phenotypic differences in egg morphology. Linear models comparing egg area–a proxy of egg size– and egg aspect ratio in six species of *Drosophila*.

| Species | df | Egg Area |  | Aspect Ratio |  |
| --- | --- | --- | --- | --- | --- |
|  |  | F | P | F | P |
| *D. melanogaster* | 18, 5178 | 76.873 | $<1 \times 10^{-10}$ | 35.474 | $<1 \times 10^{-10}$ |
| *D. santomea* | 4,1414 | 27.56 | $<1 \times 10^{-10}$ | 28.436 | $<1 \times 10^{-10}$ |
| *D. sechellia* | 5,1668 | 22.183 | $<1 \times 10^{-10}$ | 3.1564 | $7.684 \times 10^{-3}$ |
| *D. simulans* | 4,1427 | 76.773 | $<1 \times 10^{-10}$ | 30.517 | $<1 \times 10^{-10}$ |
| *D. teissieri* | 4,1445 | 142.08 | $<1 \times 10^{-10}$ | 1.0092 | 0.402 |
| *D. yakuba* | 5, 1412 | 43.824 | $<1 \times 10^{-10}$ | 25.139 | $<1 \times 10^{-10}$ |

Egg aspect ratio, a proxy of egg shape, also differed among species (LM, F_28,19866_ = 204.02, *p* < 1 × 10^-10^) and about 80% of the pairwise comparisons between species were significantly different. *Zaprionus davidi* had the highest egg aspect ratio (3.428 ± 0.739; *t* = 25.044, *p* < 1 × 10^-10^), contrasting with *Z. gabonicus* (2.268 ± 0.345; *t* =-14.457, *p* < 1 × 10^-10^), which had the lowest aspect ratio. Mean egg aspect ratio and mean egg size were not correlated at the species level (Pearson’s product-moment correlation: ρ = 0.237, *p* = 0.215).

### Egg size differs within lines of the same species

For six species, we surveyed egg traits for more than two isofemale lines per species. We fitted linear models for each of these species to assess whether egg area and aspect ratio differed within species. Egg area differed among isofemale lines for each of the six species, and aspect ratio differed in all species except for *D. teissieri* (Table 1).

We tested whether this result could be generalized. A linear model with all the six species and accounting for the species effect revealed that the line effect–nested within the species effect– was significant for both egg area (*F*_41,19825_ = 69.477, *p* < 1 × 10^-10^) and ratio aspect (*F*_41,19825_ = 22.059, *p* < 1 × 10^-10^). These results indicate that phenotypic variation within species in egg phenotypes is a common phenomenon across *Drosophila* species.

### Egg size shows negative allometry with adult body size

We measured the adult body size of males and females to calculate the power functions between egg and adult size. We found a large range of body sizes across species for both sexes. The largest species was *Zaprionus tsacasi* (mean female mass = 2.87 mg, mean male mass = 2.14 mg), and the smallest was *D. paulistorum* (mean female mass = 0.74 mg, mean male mass = 0.43 mg). Since adult mass was measured *en masse*, we did not do comparisons for this trait. Table S4 lists the estimates of the mean mass per sex for each species.

We measured the body size scaling of egg size using a suite of PGLS models and compared the estimated slopes to the null expectation of isometric scaling. Egg size was greatest, relative to adult female body mass, in *Drosophila sechellia* at 0.049 mm^2^g^-⅔^ and smallest in *D. funebris* at 0.108 mm^2^g^-⅔^ with a mean across species of 0.071 mm^2^g^-⅔^. Because egg size was measured as the area of each egg within images (as opposed to a volumetric or mass-based measurement), the expected null isometric scaling factor, as related to adult body mass (which is proportional to volume) is M_b_^0.67^. Linear measurements of egg size (major and minor axis lengths) would be expected to scale as a factor of M_b_^0.33^ with respect to adult body mass. First, we examined the power function between egg size and female body size. Egg size, measured as area, scaled relative to female body mass as M ^0.33^ (*T* = 4.51, *p* = 0.0001, Figure 2), and with male body mass as M ^0.26^ (*T* = 4.29, *p* = 0.0002). In both cases, the exponent was roughly half the expected scaling value (Table 2). Egg size (relative to female size) also scaled differently between the two major *Drosophila* clades. The *Sophophora* subgenus had a scaling factor of 0.19 ± 0.17 (Figure 2), but this was not statistically distinct from a slope of 0 (*p* = 0.282); the *Drosophila* subgenus showed a scaling factor of 0.47 ± 0.21 (*p* = 0.040).

**FIGURE 2.**
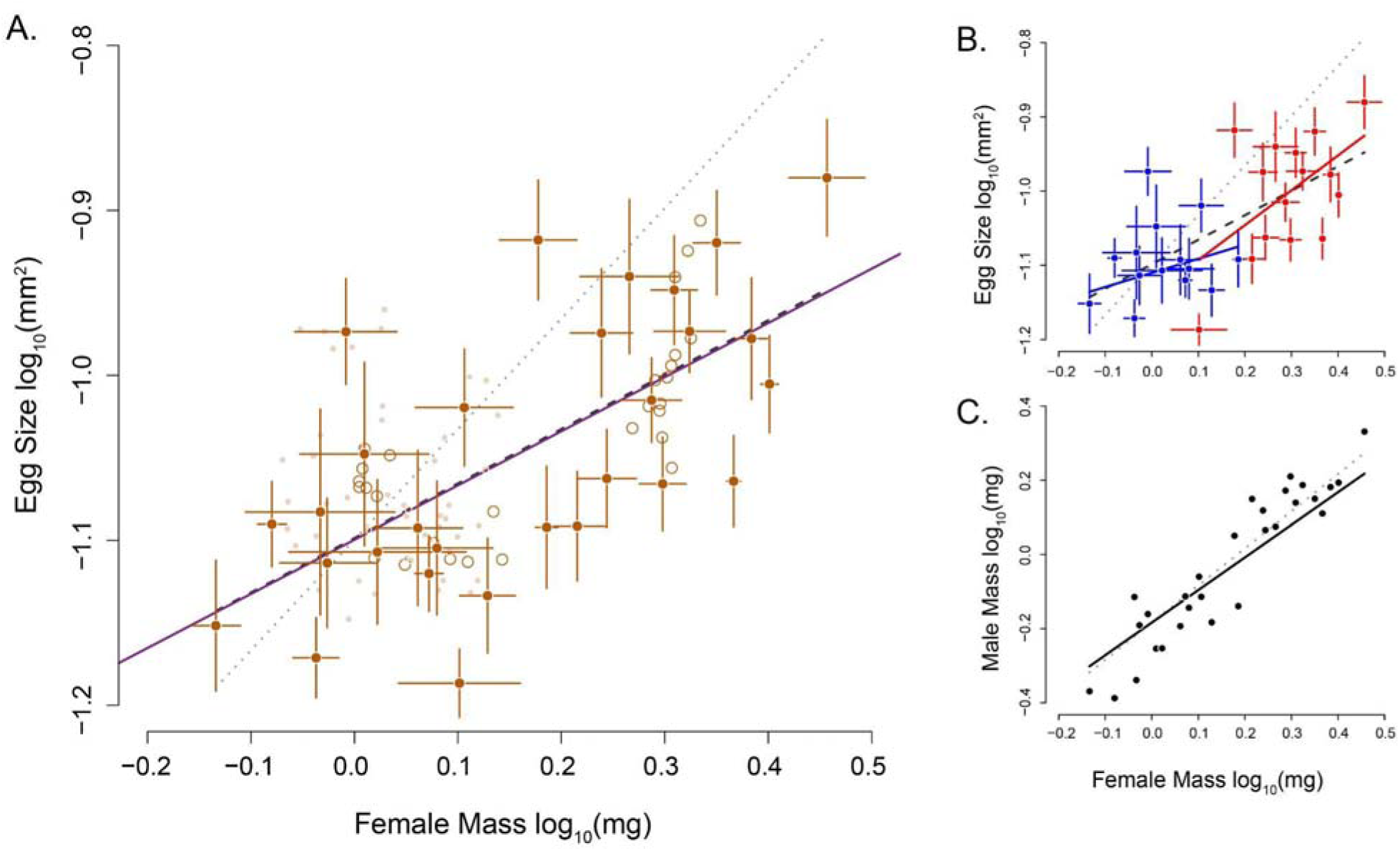
Allometry of egg size with relation to adult female body mass. **A.** Dark closed points are species means with ± 1 standard deviation error bars. Small closed points depict individual isofemale lines within some species. Open points show node values from the ancestral character state reconstruction. The results of our PGLS model are depicted by the solid violet line, and we provide two references for comparison. The slope in this log-log model represents the allometric scaling factor of egg size relative to adult body size. The dark gray dashed line shows a slope of 0.33, and the lighter grady dotted line shows the isometric null expectation (slope = 0.67). Both reference lines share their intercepts with the PGLS regression. **B.** Blue shows species means with ± 1 standard deviation error bars for the *Drosophila* subgenus and red denotes the *Sophophora* subgenus. The red and blue regression lines show the results of PGLS models for each clade compared to the isometric null (gray dotted line). **C.** Male body size vs. female body size. The solid black line shows the PGLS regression compared to a 1:1 null (gray dotted line).

**TABLE 2.** Allometric scaling relationships between egg morphology and adult mass.

| Trait | Isometry | Scaling factor* | T | P |
| --- | --- | --- | --- | --- |
| Egg Size x Female Mass | 0.67 | $0.328 \pm 0.073$ | 4.513 | 0.00011 |
| Egg Size x Male Mass | 0.67 | $0.261 \pm 0.061$ | 4.288 | 0.00021 |
| Nonphylogenetic Model | 0.67 | $0.328 \pm 0.073$ | 4.513 | 0.00011 |
| Major Axis x Female Mass | 0.33 | $0.158 \pm 0.064$ | 2.460 | 0.02058 |
| Minor Axis x. Female Mass | 0.33 | $0.146 \pm 0.040$ | 3.620 | 0.0012 |

\* Slope estimated in a log-log model ± std. error

Similarly to the area power functions, the lengths of the major and minor axes scaled at about half the expected value (0.158 and 0.146 for the major and minor axes, respectively; *p* = 0.021 and 0.001, respectively). Unlike any of the three size measurements, egg aspect ratio did not vary significantly with female body size (i.e., showed no scaling; PGLS, *F*_1,27_ = 0.254, *p* = 0.619). Since these exponents are lower than the expectation under an isometric function, these results constitute evidence of negative allometry for all the measured egg traits with the exception of aspect ratio. The implication of these results is that larger species produce smaller eggs than predicted by adult size, and that egg shape is consistent among all species in our sample.

### Phylogenetic signal

Next, we calculated the phylogenetic signal of egg size and aspect ratio, as well as adult body size. We estimated the strength of the phylogenetic signal using two complementary metrics: Pagel’s λ (Pagel, 1999) and Blomberg’s *K* (Blomberg *et al*., 2003). The λ values for all measured traits were significantly less than 1, but most were also significantly greater than 0 (Figure 3, Table 3). Egg size, female mass, and relative egg size all show moderate and significant phylogenetic signal (λ = 0.28, 0.68, and 0.41, respectively; Table 3). Unlike the other traits, egg aspect ratio showed no phylogenetic signal as its λ value was statistically indistinguishable from 0 (λ = 0.14, *p* = 0.252). Phylogenetic signal was also low within each of the two subgenera, with λ values of lambda that were similarly indistinct from 0 (λ = 0.18, *p* = 0.546 for *Sophophora*; λ = 7.34 *×* 10^-5^, *p* = 1.00 for *Drosophila*).

**FIGURE 3.**
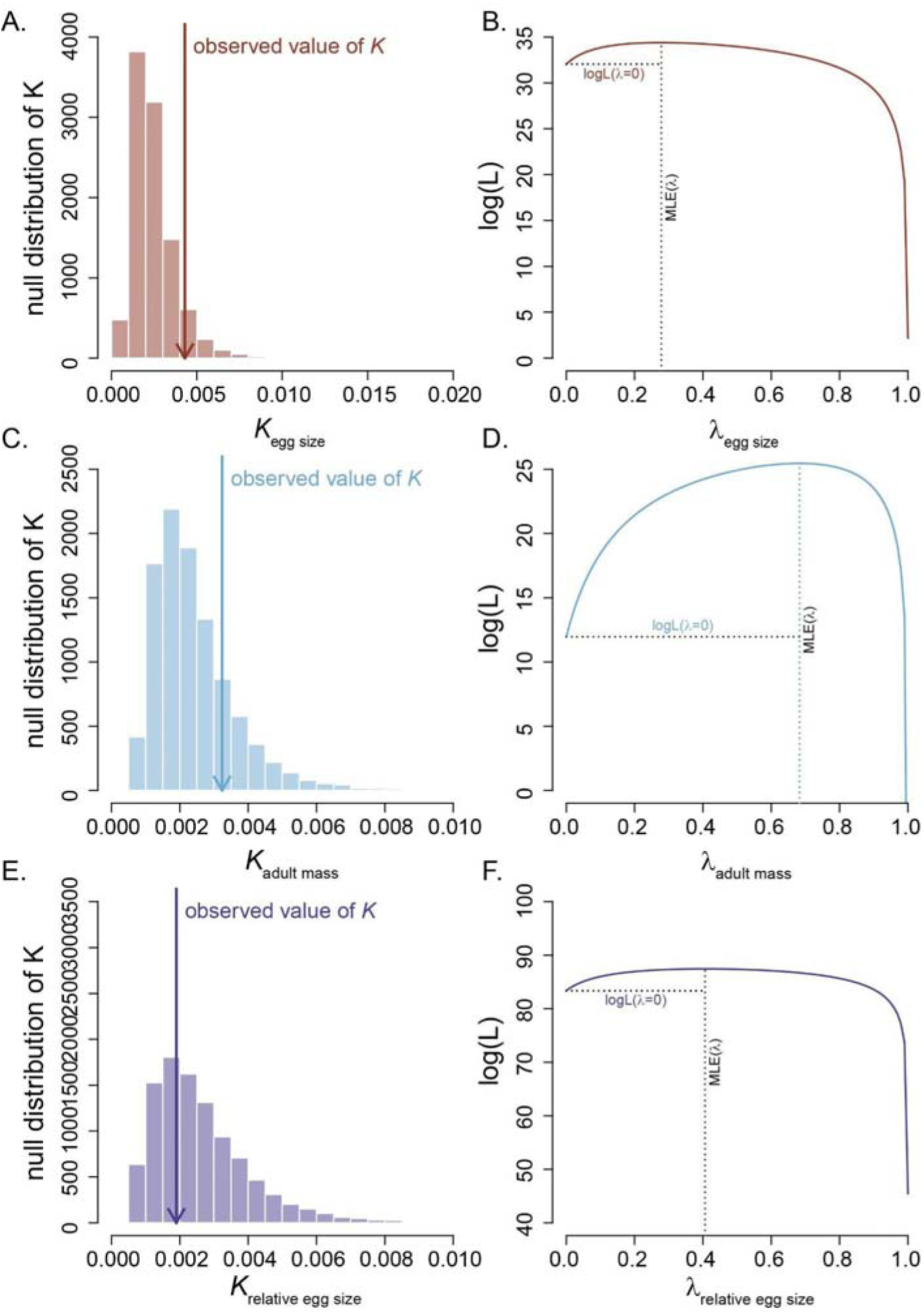
Phylogenetic signal in egg size, adult size and relative egg size in *Drosophila*. Right panels (**A.**, **C.**, and **E.**) show the distribution of Blomberg’s *K*. The left panels (**B.**, **D.**, and **F.**) show the distribution of Pagel’s λ. Top panels show analyses for egg size, middle panels show analyses for adult size, lower panels show analyses for relative egg size.

**TABLE 3.** Phylogenetic Signal for three egg morphology phenotypes and female mass.

| Trait | $\rho$ | $P_{\rho=0}$ | $P_{\rho=1}$ | $K$ | $P_{K=0}$ | $P_{K=1}$ |
| --- | --- | --- | --- | --- | --- | --- |
| Egg Size | 0.278 | 0.030 | $1.03 \times 10^{-15}$ | 0.004 | 0.092 | $9.99 \times 10^{-5}$ |
| Egg Aspect Ratio | 0.142 | 0.252 | $1.05 \times 10^{-21}$ | 0.001 | 0.832 | $9.99 \times 10^{-5}$ |
| Relative Egg Size | 0.407 | 0.004 | $5.08 \times 10^{-20}$ | 0.002 | 0.622 | $9.99 \times 10^{-5}$ |
| Female Mass | 0.683 | <0.001 | $1.16 \times 10^{-21}$ | 0.003 | 0.196 | $9.99 \times 10^{-5}$ |

We also measured Blomberg’s *K* for these traits. For all measured traits (male mass, egg size, egg/adult ratio, and female mass), Blomberg’s *K* was not significantly different from 0 (Table 4), suggesting greater trait variation than would be expected under a pure BM model of trait evolution. This result suggested that other models of trait evolution might better reflect the evolutionary dynamics of these size traits. Surprisingly, and despite the low value of Blomberg’s *K*, the best model to explain the evolution of egg size in *Drosophila* is a BM model (Table 4). We ran a similar analysis restricting the analyses to three models, WN, BM, and OU, which allowed us to more directly test the hypothesis of whether the evolution of each trait followed a BM model of trait evolution, or whether there was evidence of stabilizing selection in which selection moves egg size to a fitness optimum on an adaptive landscape (OU model). In this case, the BM model was also favored (Table S5). Regardless of the approach we use, the BM model is more likely to explain the evolution of egg size in *Drosophila* than other alternative, more complex models. We followed a similar approach to study the evolution of egg aspect ratio. The results are similar to egg size as BM is the best fitting model in the expanded set of models (Table 4). BM was the best fit for both sexes in the reduced set of models (Table S7). These results are therefore inconsistent with the existence of egg aspect ratio optima. The *Sophophora* clade showed strong support for a BM model of evolution (AIC_w_ = 0.68), but the *Drosophila* subgenus showed equivalent support for both a BM model and a non-evolutionary WN model (both AIC_w_ = 0.47).

**TABLE 4.** Evolutionary models of trait evolution in egg phenotypes. The left side of the table shows parameter estimates for aspect ratio; the right side shows parameter estimates for egg size. For both traits the BM model was the best fit.

|  | Aspect ratio |  |  |  |  | Egg size |  |  |  |  |
| --- | --- | --- | --- | --- | --- | --- | --- | --- | --- | --- |
| Model | Sig2 | SE | AICc | ΔAIC | AIC<sub>w</sub> | Sig2 | SE | AICc | ΔAIC | AIC<sub>w</sub> |
| white | 0.067 | 0 | 10.825 | 1.312 | 0.178 | 0 | 0.017 | -145.848 | 4.636 | 0.0340 |
| lambda | 0.118 | 0 | 12.220 | 2.707 | 0.089 | 0.00052 | 0 | -147.777 | 2.707 | 0.089 |
| BM | 0.017 | 0.240 | 9.513 | 0 | 0.343 | 0.00014 | 0.015 | -150.483 | 0 | 0.346 |
| OU | 0.017 | 0.240 | 12.220 | 2.707 | 0.089 | 0.00014 | 0.015 | -147.777 | 2.707 | 0.089 |
| EB | 0.083 | 0.240 | 11.960 | 2.447 | 0.101 | 0.0017 | 0.015 | -148.752 | 1.732 | 0.146 |
| rate.<br>trend | 0.051 | 0.239 | 11.911 | 2.398 | 0.103 | 0.00134 | 0.015 | -148.782 | 1.702 | 0.148 |
| delta | 0.016 | 0.241 | 12.027 | 2.514 | 0.098 | 0.00015 | 0.015 | -148.782 | 1.702 | 0.148 |

Mean body mass followed a BM model of trait evolution in females and a rate trend model in males; the support was not overwhelming in either case (Tables S6). When we used a reduced set of evolutionary models (WN, OU, and BM), the BM model was the preferred model for both female and male body size (AIC_w_=0.795 in both cases, Table S7). These last results pertaining body size should be considered preliminary and taken with caution as the metric for individual adult size was the mean of a pool of individuals.

### Evolutionary rates

Finally, we asked whether the evolutionary rate varied across the phylogeny for egg size or adult body size. We measured evolutionary rate ( ^2^) across the tree using maximum likelihood-optimized values of λ for each trait. The λ that we used for reconstructions of ^2^ for adult body size was λ *=* 0.68, while the λ for egg size was 0.27. These traits generally follow similar patterns (Figure 4). ^2^ varies tenfold for egg size and almost 1,000-fold for female adult body size. In adult body size, the ^2^ is higher in the *Drosophila* subgenus than in the *Sophophora* subgenus. Additionally, the *D. melanogaster* subgroup showed higher evolutionary rates than the *saltans* subgroup. Evolutionary rates for egg size showed greater variability (Figure 5), and the rates of evolution of the two traits were only loosely correlated across the phylogenetic tree (Pearson correlation coefficient *r* = 0.15). A noteworthy exception occurred in the *D. lummei* / *D. virilis* clade, where the evolutionary rate of adult body size was comparatively slow relative to the evolutionary rate of egg size.

**Figure 4.**
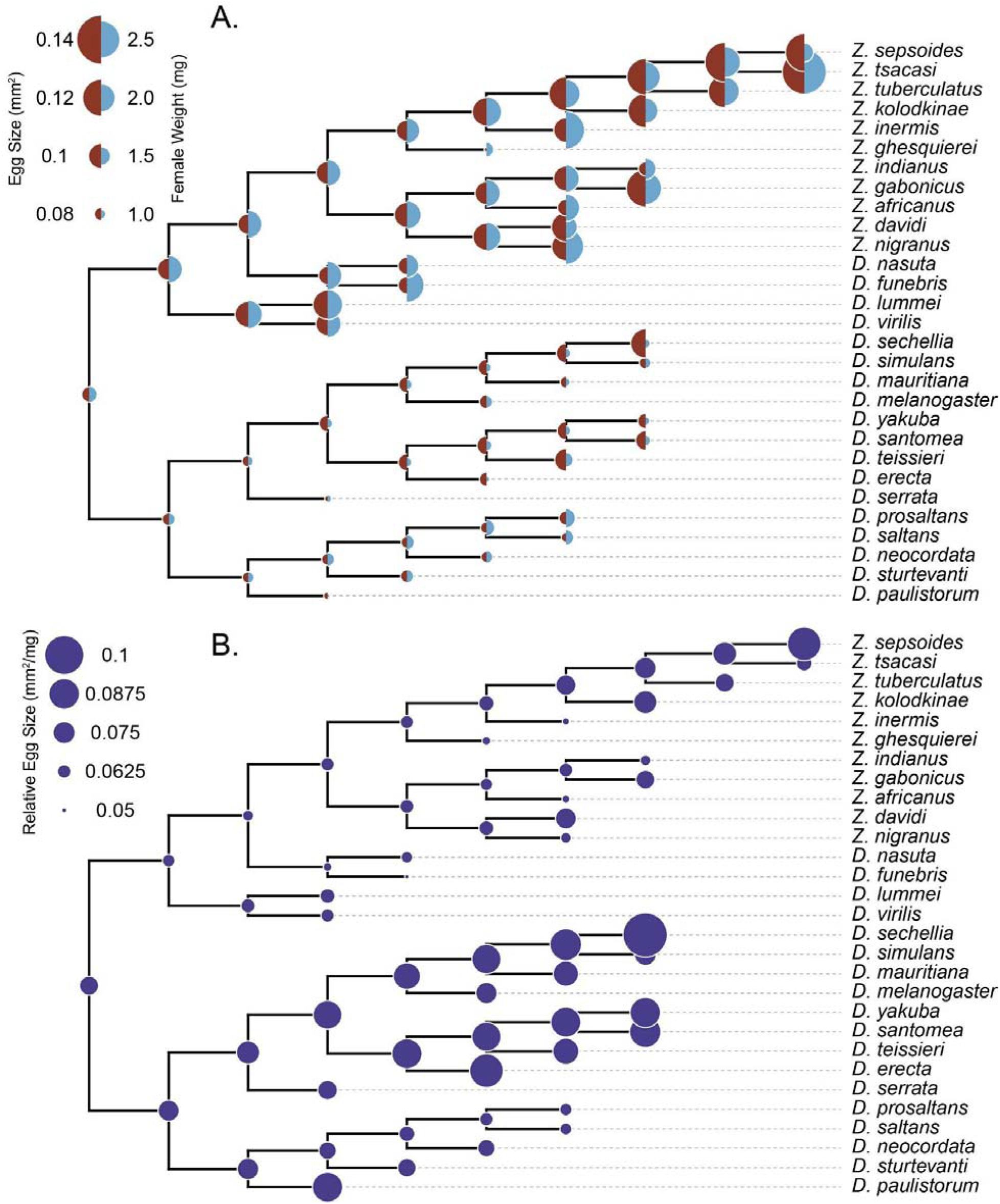
Ancestral state reconstruction. Tree branch lengths were all equalized and set to length of 1 to show tree topology. **A.** Each node shows the inferred egg size (brown, left half side) and female adult mass (blue, right half side). **B.** Relative egg size (ratio of egg size to adult female mass).

**Figure 5.**
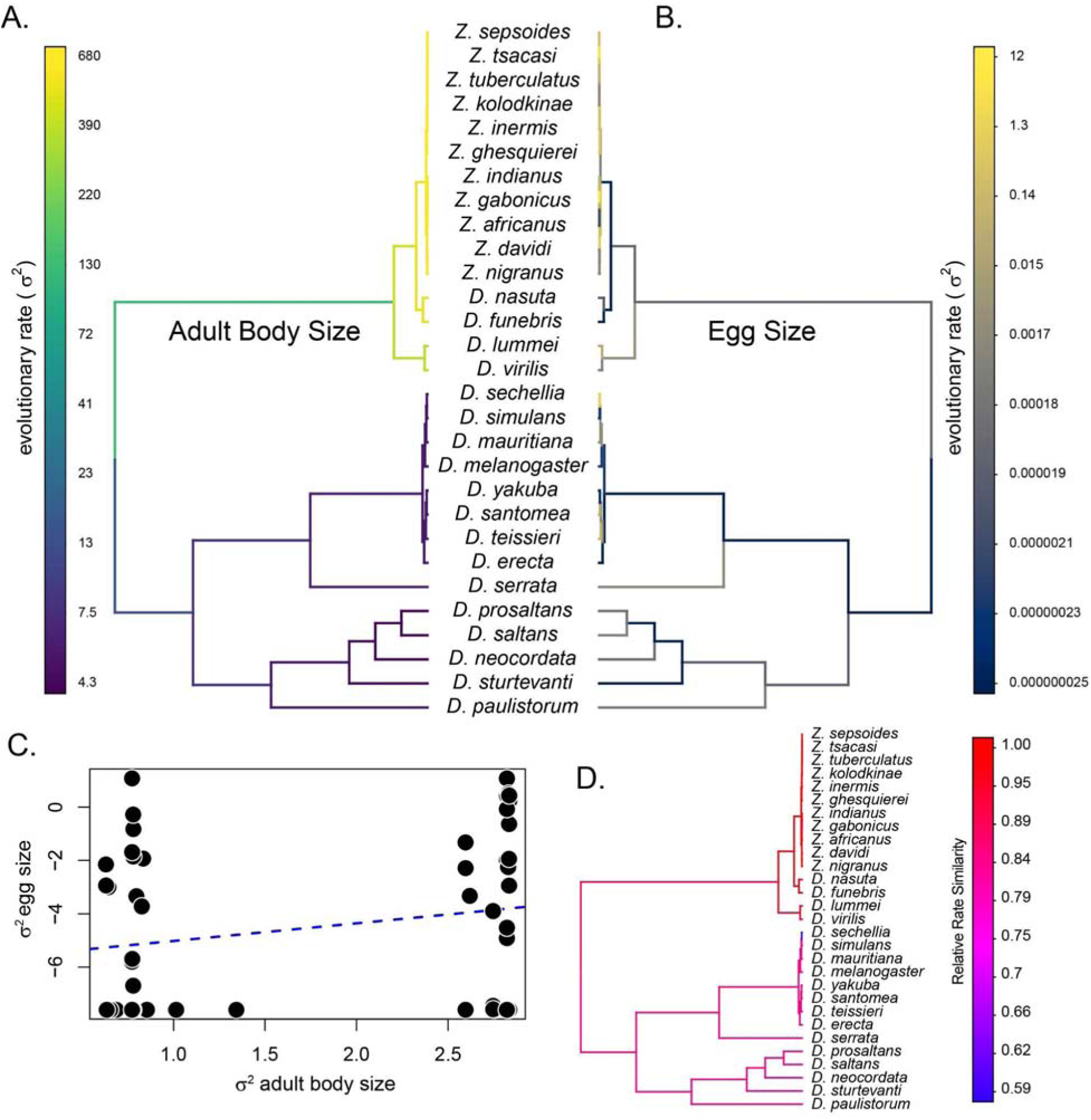
Evolutionary rate of female adult body size and egg size in drosophilids. Top two panels (**A., B.**) show the estimated rate of trait evolution (σ^2^) across the phylogeny for female adult body size and egg size. The color on the branches represents the evolutionary rate. **C.** σ^2^ values for the two traits show no significant correlation. **D.** Comparison of normalized σ^2^across the phylogeny shows that the evolutionary rate of egg size is, with a single exception, greater than the evolutionary rate of adult body size. The lone exception, shown by a blue branch, is *D. sechellia*.

## Discussion

Body size has been an active subject of research in the field of evolutionary biology since the inception of the field (Darwin & Wallace, 1858; Darwin, 1860; Alfred Wallace, 1889; Rensch, 1948). Understanding the dynamics of the trait has become a tenet of ecological studies (Peters, 1983; LaBarbera, 1989; Woodward *et al*., 2005; White *et al*., 2007). In fact, several of the known ‘rules of biology’ pertain to body size trends and species distributions (e.g., (Bergmann, 1848; Rensch, 1948; Foster, 1964). Natural selection and sexual selection often favor larger adult body size (Kingsolver & Pfennig, 2004; Hone & Benton, 2005; Kingsolver & Huey, 2008). Selection for larger adult body size is eventually counterbalanced by opposing selective forces, usually associated with delays in reproduction and increased mortality in pre-reproductive life stages due to prolonged development (reviewed in Blanckenhorn, 2000). Yet the evolution of immature body sizes remains relatively unexplored, and especially in a phylogenetically-explicit framework. In this piece, we address that gap using *Drosophila* as a model. First, we report extensive variation in egg size among species and within species. Second, we describe that egg size scales allometrically with body size in drosophilids. Finally, we describe that both, egg size and egg shape, have a moderate phylogenetic signal and follow a Brownian model of trait evolution. We elaborate on the implications of these findings in the following paragraphs.

First, we found that egg size and egg shape show an allometric relationship with adult body size, and that relative egg size decreases compared to the rest of the body in larger drosophilid species. An intriguing outcome of our study is that not only does the size of eggs not scale isometrically with respect to female body size (0.67 here, imposed by the area:volume ratio), it actually more closely approximated the expected isometric under a completely different scenario (linear:volume - 0.33). This finding may point to some limiting tradeoff in oogenesis or oviposition. For example, while the volume of eggs (and therefore its mass) may be under selection that favors at least isometric scaling with respect to adult size. All else being equal, though, the relevant measurement of an ovipositor that determines the size of eggs it can pass is its cross-sectional area, which, when you assume a circular cross-sectional shape, becomes a function of its radius - a linear metric. Furthermore, ovipositor size has a negative allometric relationship with respect to female size, at least in some species of *Drosophila* (Matsuo, 2025), though a more taxonomically-broad survey would be needed to ascertain if this is a more general trend. If the size of the ovipositor is indeed a limiting factor, we may therefore expect the scaling factors on the order of 0.33 as we have found here. It is also possible that other cryptic factors are at play governing the size of eggs.

In birds, egg size and female body size are correlated, and larger species tend to lay smaller eggs than expected given their size (Olsen *et al*., 1994; Mytiai *et al*., 2021), a pattern similar to our observations in *Drosophila*. In birds, female body weight accounts for the vast majority of variation in egg volume, with sexual dimorphism also having a moderate effect (Olsen *et al*., 1994; Weatherhead & Teather, 1994). Moreover, in birds, egg size and shape show a negative allometric relationship with pelvis size and habitat (Shatkovska *et al*., 2018). Nonetheless, analyses at finer taxonomic scales have revealed that different bird clades differ in the scaling exponents of their power functions (Olsen *et al*., 1994; Weatherhead & Teather, 1994). Of note, in species that show egg sexual size dimorphism, egg sexual and adult size dimorphism are not correlated (Rutkowska *et al*., 2014). In reptiles, egg size also shows a negative allometric relationship with female body size (Deeming & Birchard, 2007). Negative allometric relationships have also been observed in killifish (Eckerström-Liedholm *et al*., 2017); however, in this clade, species that experience more seasonal environments tend to produce larger eggs, likely reflecting increased provisioning as a bet-hedging strategy compared to species inhabiting more stable environments (Eckerström-Liedholm *et al*., 2017). This pattern is not universal. In black flies (Diptera: Simuliidae), egg size covaried with body size in only 2 of 17 surveyed species (Malmqvist *et al*., 2004). Finally, in the most comprehensive survey to date, egg size and shape in insects were found to follow no universal scaling law; instead, strong evidence supports convergent evolution in both traits driven by ecological factors (Church *et al*., 2019).

The study of allometric relationships can be extended to other developmental stages, to the rate of development and the factors that regulate it, and ultimately to gene expression. Several precedents indicate that examining allometric relationships in immature stages can reveal evolutionary patterns that govern growth rates. For example, allometric exponents in lepidopteran larvae vary among species by as much as a factor of two (Tammaru & Esperk, 2007). In aquatic insects, including vector species such as *Aedes albopictus*, length–mass allometry slopes in larval stages vary both between and within families (Mocq *et al*., 2024), with implications for ecological studies that require biomass estimates (Eklöf *et al*., 2017) as well as for vector biology (Nørgaard *et al*., 2022). The rate of size change itself is also an important question that benefits from a comparative phylogenetic perspective. For instance, incubation periods in reptiles and birds follow an allometric relationship with initial egg mass, although in birds the magnitude of this relationship is order-specific (Deeming *et al*., 2006). Gene expression analyses in three *Drosophila* species revealed interspecific differences in the spatial positioning of expression domains of developmental regulators during embryogenesis, alongside strong maternal control within *D. melanogaster* (Lott *et al*., 2007). Another promising research avenue is to understand the cellular and biophysical underpinnings of allometric relationships (Rensch, 1948; Shingleton *et al*., 2007). In Hawaiian *Drosophila*—a clade in which body length spans a fourfold range—different organs (eye, wing, and basitarsus) show negative allometric relationships with cell size (Stevenson *et al*., 1995).

A second finding of our survey is that egg traits in *Drosophila* show moderate phylogenetic signal and show patterns of evolution consistent with a Brownian motion (BM) model. The relatively low phylogenetic signal indicates that egg size and morphology in *Drosophila* display limited phylogenetic conservatism. These results contrast with the strong phylogenetic signal of egg size reported in birds (Shatkovska *et al*., 2018), and cephalopods (Ibáñez *et al*., 2021). Similarly, our finding that the BM model provides the best fit in *Drosophila* contrasts with results from birds (Cooney *et al*., 2020), cephalods, and reptiles (Oliveira *et al*., 2025), none of which are best explained by a BM model. Notably, analyses across insects as a whole indicate that egg width and egg length show strong phylogenetic signal and that an OU model provides the best fit (Soulsbury & Iossa, 2024). Together, these differences among taxa indicate that the evolutionary tempo and mode of egg size and shape vary markedly across groups.

Gamete and embryo size are core components of life-history theory. A classic life-history trade-off predicts that species laying larger eggs should also produce fewer offspring. In drosophilids, egg size and ovariole number are negatively correlated, even after accounting for adult body size. This negative correlation has been documented in Hawaiian *Drosophila* (Montague *et al*., 1981), and both traits are also influenced by oviposition substrate (Sarikaya *et al*., 2019). However, the relationship between egg size and reproductive output is not universal across insects (Church *et al*., 2021), and holds only at a qualitative level in lizards (Sinervo *et al*., 1992). Indeed, this pattern is absent in several insect clades (e.g., Orthoptera, Coleoptera, and Hymenoptera), which show no significant correlation between egg size and the number of eggs laid per female, a pattern that is not affected by parental care mode (Gilbert & Manica, 2010). A second proposed trade-off is that larger eggs develop faster because they are better provisioned than smaller eggs, a prediction supported by several theoretical models. For example, in birds, initial egg size shows an almost isometric relationship with hatchling mass (Olsen *et al*., 1994). *Drosophila*, however, does not follow this pattern (Markow *et al*., 2009). More broadly, clades with postnatal provisioning (birds and mammals) exhibit stronger correlated evolution between juvenile mass at independence and adult body size than clades lacking postnatal provisioning (fish, anurans, and reptiles; Rollinson *et al*., 2019). Nonetheless, larger eggs and larger juveniles do not always confer the highest survival. In *Drosophila*, a survey of three tropical species suggested that female body size does not predict ovariole number (Krishna *et al*., 2012). To date, no study has systematically examined the integration between adult body size, egg number per female, and egg size in *Drosophila*. Similarly, power-function relationships across larval instars have not been studied in *Drosophila* or, more generally, in insects. The coevolution of phenotypes and their integration across life stages, therefore, remains an important area of biology.

## Author contributions

JAR and DRM conceived of the project

JAR and DRM designed the study

MP and DAC collected data JAR conducted analyses

JAR, MP, and DRM wrote the initial manuscript draft

All authors edited the manuscript

## Acknowledgements

We thank the Matute lab for their helpful comments on this manuscript. Funding for this work was provided by the National Institute of Environmental Health Services (F32ES035271) to Jonathan A. Rader, and the National Institute of General Medical Sciences (R35GM148244) to Daniel R. Matute.

## Data Accessibility Statement

The raw and processed egg images, along with our dataset of egg measurements and analysis scripts (Fiji and R) are archived at FIGSHARE.

## Supplementary Figures

**Figure S1.**
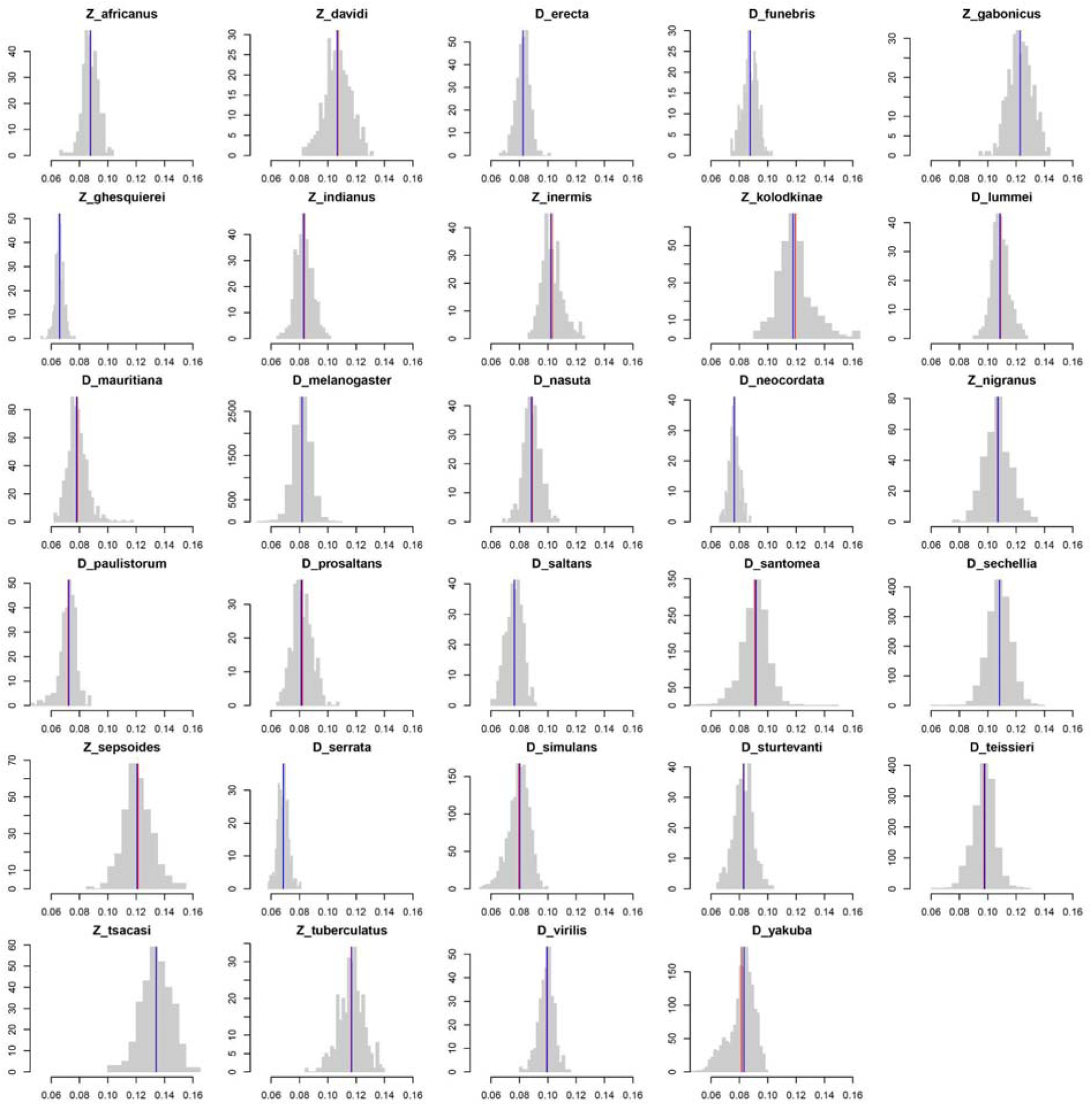
Histograms of egg size by species. Common x-axis range allows comparison across taxa. Red vertical line shows species mean, blue vertical line shows species median.

## Supplementary Tables

**TABLE S1.** Stocks used in this study.

| Species | Isofemale line | Location | Collector |
| --- | --- | --- | --- |
| *Z. africanus* | 18. KARI. 21. Z. afr | Kenya | D. Matute |
| *Z. inermis* | 18. TRNE. 05. Z04. Z. inerm | Sao Tóme | D. Matute |
| *Z. nigranus* | 18CAR07. Z03. Z. nig | Sao Tóme | D. Matute |
| *Z. tuberculatus* | 18was04. Z. tub | Sao Tóme | D. Matute |
| *D. nasuta* | 18was14. D. nasuta | Stock Center |  |
| *D. sechellia* | C24 | Seychelles |  |
| *D. sechellia* | C38 | Seychelles |  |
| *D. sechellia* | C42 | Seychelles |  |
| *D. sechellia* | C46 | Seychelles |  |
| *D. sechellia* | C50 | Seychelles |  |
| *D. sechellia* | C54 | Seychelles |  |
| *D. erecta* | D. erecta | Stock Center |  |
| *D. mauritiana* | D. mau12 | Mauritius | Unknown |
| *D. serrata* | D. serrata. light. line1 | Stock Center |  |
| *D. funebris* | Funebris. 0. 01 | Stock Center |  |
| *D. lummei* | Lummei. 15010. 1011. 01 | Stock Center |  |
| *D. melanogaster* | LUN15. 025 | Zambia | D. Matute |
| *D. melanogaster* | LUN15. 045 | Zambia | D. Matute |
| *D. melanogaster* | LUN15. 051 | Zambia | D. Matute |
| *D. melanogaster* | mauritiana | Mauritius | Unknown |
| *D. melanogaster* | MEL. LZV. L36 | Zambia | D. Matute |
| *D. melanogaster* | MEL. LZV. L36. 2 | Zambia | D. Matute |
| *D. melanogaster* | MEL. LZV. L43 | Zambia | D. Matute |
| *D. melanogaster* | MEL. LZV. L43. 2 | Zambia | D. Matute |
| *D. melanogaster* | MEL. LZV. L69 | Zambia | D. Matute |
| *D. melanogaster* | MEL. LZV. L69. 2 | Zambia | D. Matute |
| *D. melanogaster* | MEL. LZV. L91 | Zambia | D. Matute |
| *D. melanogaster* | MEL. LZV. L91. 2 | Zambia | D. Matute |
| *D. melanogaster* | MEL. LZV. L93 | Zambia | D. Matute |
| *D. melanogaster* | MEL. LZV. L93. 2 | Zambia | D. Matute |
| *D. melanogaster* | Melano. DRM81. LIV | Zambia | D. Matute |
| *D. melanogaster* | Melano. DRM90. LIV | Zambia | D. Matute |
| *D. melanogaster* | Melano. DRM96. LIV | Zambia | D. Matute |
| *D. neocordata* | Neocordata. 14041. 0831. 00 | Stock Center |  |
| *D. melanogaster* | NMB15. 014 | Namibia | D. Matute |
| *D. melanogaster* | NMB15. 028 | Namibia | D. Matute |
| *D. melanogaster* | NMB15. 029 | Namibia | D. Matute |
| *D. paulistorum* | Paulistorum. 06 | Stock Center |  |
| *D. prosaltans* | Prosaltans. 14045. 0911. 00 | Stock Center |  |
| *D. santomea* | S2 | Sao Tóme | D. Matute |
| *D. santomea* | S32 | Sao Tóme | D. Matute |
| *D. santomea* | S38 | Sao Tóme | D. Matute |
| *D. saltans* | Saltans. 14045. 0911. 02 | Stock Center |  |
| *D. santomea* | San. 18. cmn. 10. 06 | Sao Tóme | D. Matute |
| *D. santomea* | San. 18. TRNE | Sao Tóme | D. Matute |
| *D. simulans* | Sim. md4 | Madagascar | Unknown |
| *D. simulans* | Sim. md41 | Madagascar | Unknown |
| *D. simulans* | Sim. N31 | Kenya | P. Andolfatto |
| *D. simulans* | Simulans. Ya20 | Cameroon | P. Andolfatto |
| *D. simulans* | Simulans. Ya9 | Cameroon | P. Andolfatto |
| *D. sturtevantii* | Sturtevantii. 14043. 0071. 21 | Stock Center |  |
| *D. teissieri* | T1 | Bioko, Equatorial Guinea | D. Matute |
| *D. teissieri* | T13 | Bioko, Equatorial Guinea | D. Matute |
| *D. teissieri* | T14 | Bioko, Equatorial Guinea | D. Matute |
| *D. teissieri* | T22 | Bioko, Equatorial Guinea | D. Matute |
| *D. teissieri* | T6 | Bioko, Equatorial Guinea | D. Matute |
| *D. virilis* | Virilis. 15010. 1051. 49 | Stock Center |  |
| *D. yakuba* | Y46 | Sao Tóme | D. Matute |
| *D. yakuba* | Yak. 1. 6 | Sao Tóme | D. Matute |
| *D. yakuba* | Yak. 1. 7. 1 | Sao Tóme | D. Matute |
| *D. yakuba* | Yak. 3. 4. 1 | Sao Tóme | D. Matute |
| *D. yakuba* | Yak. cameroon | Cameroon | Unknown |
| *D. yakuba* | Yak. Y | Cameroon | Unknown |
| *Z. davidi* | Z. davidi. JD18 | Sao Tóme | J. David |
| *Z. gabonicus* | Z. gabonicus. JD18 |  | J. David |
| *Z. ghesquierei* | Z. ghesquierei. SEN18. SYN12.<br>Pg1N | Senegal | D. Matute |
| *Z. indianus* | Z. ind. rcr. 17. 06 | Senegal | D. Matute |
| *Z. kolodkinae* | Z. kolodkinae. JD18 |  | J. David |
| *Z. sepsoides* | Z. sepsoides. Mayott. JD18 | Mayotte | J. David |
| *Z. tsacasi* | Z. tsacasi. ST. 16. 4 | Sao Tóme | D. Matute |

**Table S2.** Species egg sample sizes.

| **Species** | **n** |
| --- | --- |
| *D. erecta* | 303 |
| *D. funebris* | 305 |
| *D. lummei* | 301 |
| *D. mauritiana* | 618 |
| *D. melanogaster* | 5197 |
| *D. nasuta* | 301 |
| *D. neocordata* | 299 |
| *D. paulistorum* | 303 |
| *D. prosaltans* | 301 |
| *D. saltans* | 304 |
| *D. santomea* | 1419 |
| *D. sechellia* | 1674 |
| *D. serrata* | 302 |
| *D. simulans* | 1432 |
| *D. sturtevantii* | 310 |
| *D. teissieri* | 1450 |
| *D. virilis* | 301 |
| *D. yakuba* | 1418 |
| *Z. africanus* | 306 |
| *Z. davidi* | 300 |
| *Z. gabonicus* | 316 |
| *Z. ghesquierei* | 307 |
| *Z. indianus* | 316 |
| *Z. inermis* | 301 |
| *Z. kolodkinae* | 301 |
| *Z. nigranus* | 314 |
| *Z. sepsoides* | 305 |
| *Z. tsacasi* | 301 |
| *Z. tuberculatus* | 290 |

**TABLE S3.** Mean mass per sex for 29 drosophilid species. Note that these estimates were derived as the population mean from at least 54 individuals.

| Species | Mean F Mass ± SD (mg) | n |
| --- | --- | --- |
| *D. erecta* | 0.832 ± 0.026 | 126 |
| *D. funebris* | 2.326 ± 0.036 | 73 |
| *D. lummei* | 2.113 ± 0.164 | 149 |
| *D. mauritiana* | 0.945 ± 0.095 | 244 |
| *D. melanogaster* | 1.157 ± 0.112 | 4446 |
| *D. nasuta* | 1.756 ± 0.116 | 125 |
| *D. neocordata* | 1.181 ± 0.037 | 135 |
| *D. paulistorum* | 0.735 ± 0.04 | 127 |
| *D. prosaltans* | 1.535 ± 0.038 | 139 |
| *D. saltans* | 1.347 ± 0.084 | 136 |
| *D. santomea* | 1.032 ± 0.152 | 679 |
| *D. sechellia* | 0.987 ± 0.106 | 1144 |
| *D. serrata* | 0.919 ± 0.047 | 109 |
| *D. simulans* | 1.072 ± 0.209 | 764 |
| *D. sturtevantii* | 1.209 ± 0.151 | 155 |
| *D. teissieri* | 1.284 ± 0.13 | 628 |
| *D. virilis* | 1.94 ± 0.129 | 147 |
| *D. yakuba* | 0.939 ± 0.153 | 863 |
| *Z. africanus* | 1.988 ± 0.104 | 188 |
| *Z. davidi* | 1.736 ± 0.121 | 117 |
| *Z. gabonicus* | 2.242 ± 0.115 | 54 |
| *Z. ghesquierei* | 1.271 ± 0.172 | 94 |
| *Z. indianus* | 1.644 ± 0.101 | 117 |
| *Z. inermis* | 2.519 ± 0.049 | 118 |
| *Z. kolodkinae* | 1.852 ± 0.208 | 61 |
| *Z. nigranus* | 2.421 ± 0.085 | 82 |
| *Z. sepsoides* | 1.509 ± 0.127 | 131 |
| *Z. tsacasi* | 2.87 ± 0.232 | 165 |
| *Z. tuberculatus* | 2.04 ± 0.103 | 275 |

**TABLE S4.** Models of trait evolution tested in this study.

| Model | Model description | $\sigma^2$ | $Z_0$ | Additional parameters |
| --- | --- | --- | --- | --- |
| White noise | Trait values are modeled as part of a normal distribution, assuming no covariance among species. | rate of evolution of a continuous trait | value for the root state | NA |
| BM | The correlation between trait values increases with the amount of shared evolutionary history between pairs of tips (Felsenstein, 1973). | idem | idem | NA |
| OU | Trait evolution is modeled as a random walk with central tendency, where the parameter $\alpha$ regulates the strength of attraction toward the optimum and varies from approximately 0 to 2.72 (Butler and King, 2004). | | | |
| EB | The rate of evolution is assumed to follow an exponential function of time, $r_t = r_0 \times e^{(a \times t)}$ | | | $r_0$: initial rate, $a$: the rate of change, $t$: time. |
| Kappa | Trait divergence is associated with splitting events in the | | | $\kappa$: exponential coefficient that determines the |
|  | phylogeny (Pagel 1999). |  |  | extent of gradual evolution |
| Lambda | Trait evolution is modeled under the $\lambda$ model, in which the correlation between species' traits is rescaled according to the parameter $\lambda$, ranging from 0 (no phylogenetic signal) to 1 (full phylogenetic signal). | | | |
| Rate trend | Trait evolution is modeled under a rate trend model, in which the evolutionary rate changes linearly or exponentially over time, allowing for acceleration or deceleration of trait evolution. |  |  |  |

**TABLE S5.** Three evolutionary models of trait evolution in average adult mass. The left side of the table shows parameter estimates for average female mass; the right side shows parameter estimates for average male mass. For both traits the BM model was the best fit.

|  | Aspect ratio |  |  |  |  | Egg size |  |  |  |  |
| --- | --- | --- | --- | --- | --- | --- | --- | --- | --- | --- |
| Model | $\sigma^2$ | SE | AICc | ΔAIC | AIC<sub>w</sub> | $\sigma^2$ | SE | AICc | ΔAIC | AIC<sub>w</sub> |
| white | 0.067 | 0 | 10.825 | 1.312 | 0.292 | 0 | 0.017 | -145.848 | 1.312 | 0.292 |
| BM | 0.017 | 0.240 | 9.513 | 0.000 | 0.563 | $1.4 \times 10^{-4}$ | 0.015 | -150.483 | 0.000 | 0.563 |
| OU | 0.017 | 0.240 | 12.220 | 2.707 | 0.145 | $1.4 \times 10^{-4}$ | 0.015 | -147.777 | 2.707 | 0.145 |

**TABLE S6.**
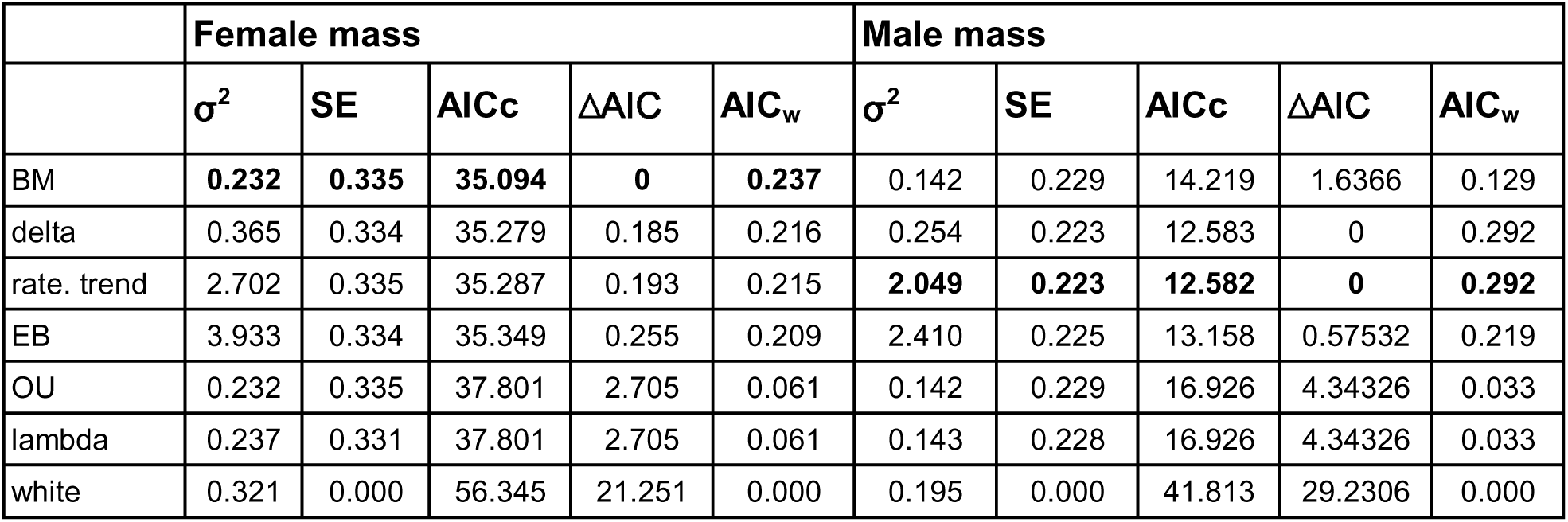
Evolutionary models of trait evolution in average adult mass. The left side of the table shows parameter estimates for average female mass; the right side shows parameter estimates for average male mass. For both traits the BM model was the best fit.

**TABLE S7.**
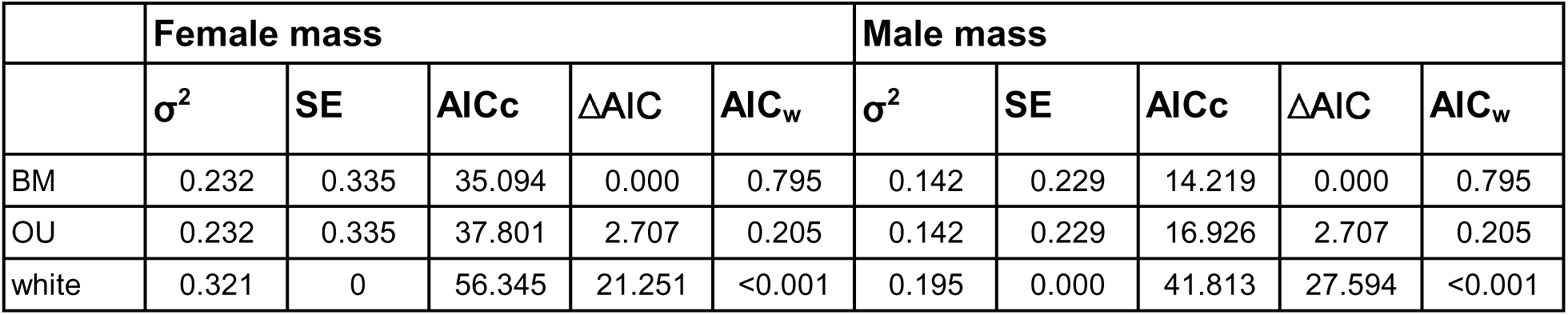
Three evolutionary models of trait evolution in average adult mass. The left side of the table shows parameter estimates for average female mass; the right side shows parameter estimates for average male mass. For both traits the BM model was the best fit.

|  | Female mass |  |  |  |  | Male mass |  |  |  |  |
| --- | --- | --- | --- | --- | --- | --- | --- | --- | --- | --- |
| | $\sigma^2$ | SE | AICc | ΔAIC | AIC<sub>w</sub> | $\sigma^2$ | SE | AICc | ΔAIC | AIC<sub>w</sub> |
| BM | 0.232 | 0.335 | 35.094 | 0.000 | 0.795 | 0.142 | 0.229 | 14.219 | 0.000 | 0.795 |
| OU | 0.232 | 0.335 | 37.801 | 2.707 | 0.205 | 0.142 | 0.229 | 16.926 | 2.707 | 0.205 |
| white | 0.321 | 0 | 56.345 | 21.251 | <0.001 | 0.195 | 0.000 | 41.813 | 27.594 | <0.001 |

**Table S8.** Evolutionary model results for relative egg size.

| | $\sigma^2$ | SE | AICc | ΔAIC | AIC<sub>w</sub> |
| --- | --- | --- | --- | --- | --- |
| BM | 0.00013 | 0.01052 | -167.952 | 0 | 0.41861 |
| delta | 0.00011 | 0.01045 | -165.482 | 2.47018 | 0.12173 |
| rate. trend | 0 | 0.01046 | -165.47 | 2.48136 | 0.12106 |
| OU | 0.00026 | 0.01047 | -165.369 | 2.58281 | 0.11507 |
| EB | 0.00013 | 0.01052 | -165.245 | 2.70667 | 0.10816 |
| lambda | 0.00016 | 0.00979 | -165.245 | 2.70667 | 0.10816 |
| white | 0.00019 | 0 | -159.83 | 8.12144 | 0.00722 |

